# Effect of sunlight on the efficacy of commercial antibiotics used in agriculture

**DOI:** 10.1101/2020.07.10.197848

**Authors:** Sebastian Khan, Amanda Osborn, Prahathees J. Eswara

## Abstract

Antibiotic stewardship is of paramount importance to limit the emergence of antibiotic-resistant bacteria in not only hospital settings, but also in animal husbandry, aquaculture, and agricultural sectors. Currently, large quantities of antibiotics are applied to treat agricultural diseases like citrus greening disease (CGD). The two commonly used antibiotics approved for this purpose are streptomycin and oxytetracycline. Although investigations are ongoing to understand how efficient this process is to control the spread of CGD, to our knowledge, there have been no studies that evaluate the effect of environmental factors such as sunlight on the efficacy of the above-mentioned antibiotics. We conducted a simple disc-diffusion assay to study the efficacy of streptomycin and oxytetracycline after exposure to sunlight for 7- or 14-day periods using *Escherichia coli* and *Bacillus subtilis* as the representative strains of Gram-negative and Gram-positive organisms respectively. Freshly prepared discs and discs stored in the dark for 7 or 14 days served as our controls. We show that the antibiotic potential of oxytetracycline exposed to sunlight dramatically decreases over the course of 14 days against both *E. coli* and *B. subtilis*. However, the effectiveness of streptomycin was only moderately impacted by sunlight. It is important to note that antibiotics that last longer in the environment may play a deleterious role in the rise and spread of antibiotic-resistant bacteria. Further studies are needed to substantively analyze the safety and efficacy of antibiotics used for broader environmental applications.

**IMPORTANCE:** Although antibiotics have been used for agricultural purposes for decades, due to the rapid rise in antibiotic resistance this usage needs to be revisited. Questions remain on the appropriate mode of application of antibiotics and the actual benefits of using antibiotics for treating the infections caused by plant pathogens, especially for the ones that are intracellular in nature. Here we show that the two commonly used commercial antibiotics, oxytetracycline and streptomycin, lose their efficacy at different rates in the presence of sunlight. While the former loses its potency within days the latter remains active for many days. Thus, oxytetracycline may not be active long enough to produce desired effect and streptomycin may persist in the environment and as a side effect due to its selective pressure, may force the rise of streptomycin-resistant pathogens.

## INTRODUCTION

Antibiotic resistance-related mortalities are expected to exceed the other leading causes of death such as cancer worldwide by 2050 [1]. Antibiotic stewardship is therefore promoted in all sectors including human health, animal husbandry, and agriculture [2–4]. The World Health Organization and the United States Centers for Disease Control and Prevention have recognized antimicrobial resistance as an enormous ongoing threat to public health [5, 6]. Runoff of antibiotics in hospital waste water [7] and intentional use in aquaculture [8], animal husbandry [9–11], and crop management [12] contribute to the rise and spread of antibiotic resistant bacteria. In this context, alarm was raised recently regarding the spraying of antibiotics in open fields as an infection control strategy to stem the spread of bacterial disease in plants [13, 14]. Specifically, the strategy approved by the United States Environmental Protection Agency [13, 15, 16] is to use streptomycin and oxytetracycline to control the spread of citrus greening disease (CGD), also known as huanglongbing (yellow dragon disease). CGD is a devastating bacterial disease caused by *Candidatus Liberibacter asiaticus* (CLas) that is transmitted between plants by certain psyllids, which are sap-feeding insects. CLas is a fastidious, Gram-negative, intracellular plant pathogen that belongs to the phylum of α-proteobacteria [17, 18]. Streptomycin and oxytetracycline are also used to treat infections caused by another bacterial plant pathogen, *Erwinia amylovora*, which causes fire blight in apples, pears, and other related species [19]. *E. amylovora* has dual growth modes - an epiphytic mode that is readily accessible for external antibiotics and an endophytic mode that is less accessible to external antibiotics [19]. In addition, tetracycline antibiotics including oxytetracycline are used in animal husbandry [20] and aquaculture [21]. Apart from the uses described above, data also suggests that antibiotics may find their way into and possibly persist in different animal and plant tissues [22–25], which could be an alternate pathway that can lead to the development of antibiotic-resistant bacteria. Thus, a comprehensive knowledge of the fate of antibiotics used in agriculture is urgently needed to hopefully curb the rise and spread of antibiotic resistance.

Although the application of antibiotics to treat CGD inspired us to pursue this study, the primary objective of this report is to investigate the effect of environmental factors, specifically sunlight, on streptomycin and oxytetracycline. To this end, we conducted a disc-diffusion assay with Gram-negative *Escherichia coli* and Gram-positive *Bacillus subtilis* and monitored the zones of inhibition of antibiotic-containing discs that were exposed to sunlight for a 7- or 14-day period. Discs that were kept in the dark for equivalent duration or that were freshly prepared served as our controls. Based on our results, we report that sunlight significantly impairs the efficacy of oxytetracycline, but only moderately impacts streptomycin. While short-lived antibiotics may not be active long enough for their intended purpose, stable antibiotics may apply constant selection pressure and create an environment conducive for the emergence of antibiotic-resistant strains [26]. Although this study (designed for undergraduate-level students [27]) is not comprehensive, our data provides a window into the life span of commercial antibiotics in nature that we hope highlights the need for further rigorous safety and efficacy investigations for the environmental use of antibiotics.

## RESULTS

### Oxytetracycline loses its antibiotic potential in the presence of sunlight in the span of few days

To monitor the effect of sunlight on the efficacy of oxytetracycline, we conducted a disc-diffusion assay. Briefly, we prepared multiple discs with oxytetracycline (50 μg) dissolved in water and placed the antibiotic-laden discs in either a natural outdoor setting with abundant sunlight to simulate agricultural use, or in a dark indoor cabinet for 7 or 14 days. In addition to the discs that were kept in the dark, we also used freshly prepared discs and vehicle (water) discs as controls. The discs were then placed, as shown in **Fig. 1**, on a pre-inoculated plate containing either a lawn of *E. coli* or *B. subtilis* cells. In all cases, as expected, the blank disc (N; negative control) and the freshly prepared discs (P; positive control) showed negligible and maximum zones of inhibition (ZOI), respectively (**Figs. 1A-D**). The discs that were kept in the dark (labeled “D”) for the duration of 7 or 14 days appeared to produce similar ZOI as our positive control of approximately 9 mm for *E. coli* and 8 mm for *B. subtilis* (**Figs. 1EF**). This suggests that oxytetracycline maintains its efficiency in the dark at room temperature for at least the maximum duration of this experiment (14 days). Next, we quantified the ZOI of the discs that were exposed to sunlight (labeled “L”) for either a 7- or 14-day period. We observed that the efficacy of oxytetracycline gradually and significantly decreased over time to almost similar to our negative control in both *E. coli* and *B. subtilis* and only retained less than 15% activity after 14 days (**Figs. 1A-F**). This implies that in the presence of sunlight, oxytetracycline loses its antibiotic potential in a matter of few days.

**Figure 1.**
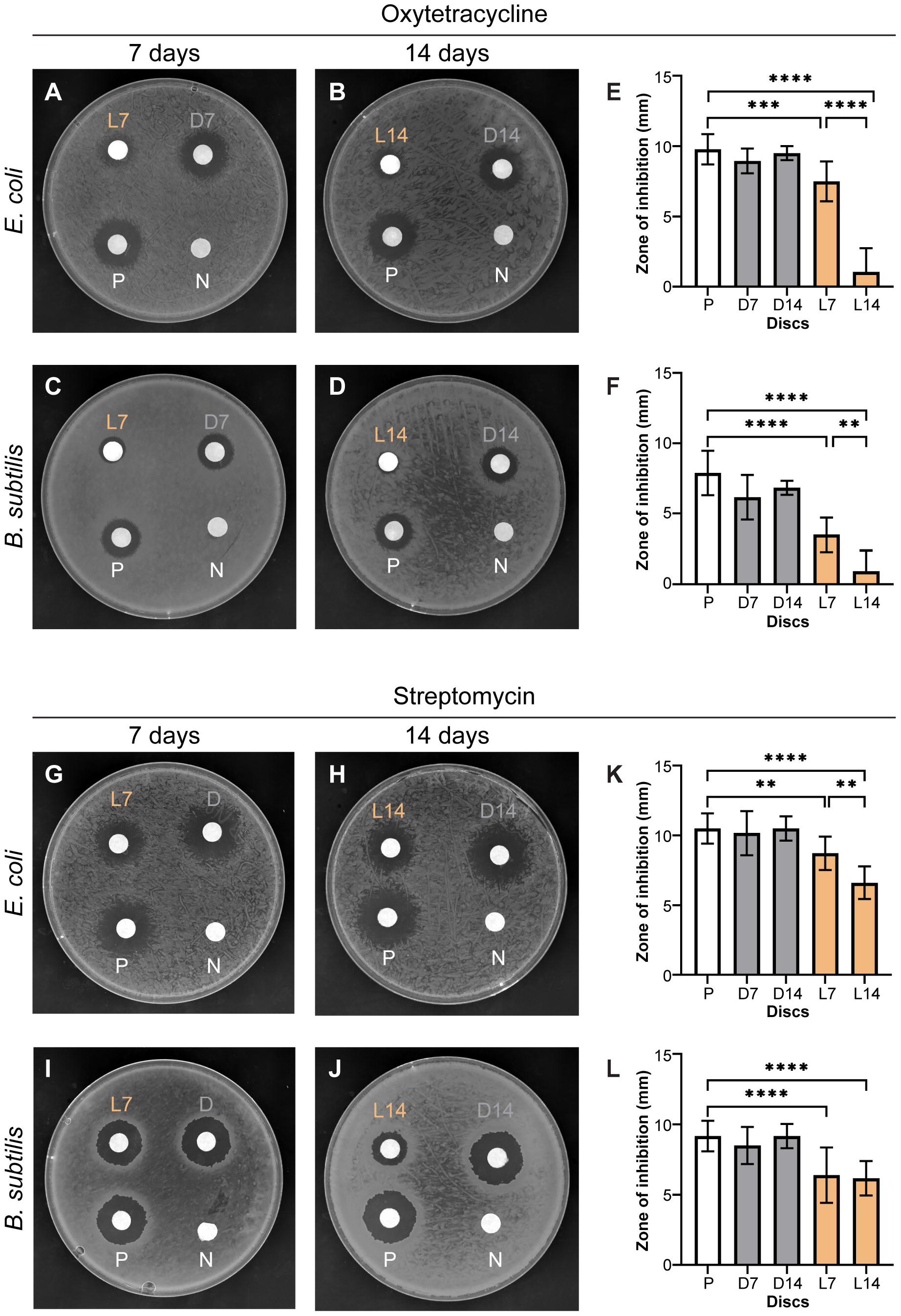
Oxytetracycline and streptomycin lose antibiotic potential in the presence of sunlight. Shown are representative disc-diffusion assay results for the effects of oxytetracycline **(A-D)** or streptomycin **(G-J)** on growth of either Gram-positive *B. subtilis* or Gram-negative *E. coli*. Quantification of the zones of inhibition in millimeters are plotted for each 7- or 14-day cohort of oxytetracycline **(E-F)** and streptomycin **(K-L).** Significance was determined using a one-way ANOVA with Tukey’s multiple comparisons analysis. Error bars represent standard deviation (SD) of the mean from three biological replicates. N: negative control (discs prepared with sterile water), P: positive control (discs prepared the day of testing), L7 or L14: 7 or 14 days in sunlight, D7 or D14: 7 or 14 days in darkness. ****: p<0.0001, ***: p<0.001, **: p<0.01.

### Moderate negative effects of sunlight on the efficacy of streptomycin

A similar experimental setup to the one discussed above was adopted for studying the effects of sunlight on streptomycin. As noted earlier, blank discs and freshly prepared discs with streptomycin (200 μg) served as our negative and positive controls respectively. As expected, the ZOI were unobservable for our blank discs and at a maximum for our positive controls (**Figs. 1G-L**). Similar to oxytetracycline, streptomycin is also able to maintain its efficacy when kept in darkness for the duration of our experiment (**Figs. 1G-L**). However, unlike oxytetracycline, streptomycin appears to be moderately resistant to sunlight. At the 7-day mark, based on the ZOI (**Figs. 1KL**), the discs exposed to sunlight appear to have retained almost approximately 80% and 70% of their activity in *E.coli* and *B. subtilis* respectively, when compared to that of our positive control. Further measurable decrease to nearly 50% efficiency compared to our positive control was noted subsequent to 14 days of sunlight exposure for *E. coli*. However, the decrease in efficiency for *B. subtilis* at the 14-day time point was within the standard error when compared to that of the 7-day time point (**Figs. 1HJKL**).

## DISCUSSION

Rapid rise of antibiotic resistance in bacteria is a major concern worldwide with enormous predicted fatalities. Antibiotics are now routinely used in clinics, animal husbandry, and agriculture. Acknowledgement of the fact that the rise of antibiotic resistance stemming from one of those settings could potentially render antibiotics useless and lead to the formation of a multidisciplinary collaborative initiative under the umbrella term One Health [2, 3]. Despite this, environmental antibiotic pollution is a growing concern that requires urgent attention [28].

Some commercial antibiotics such as oxytetracycline and streptomycin are produced by soil-dwelling *Streptomyces* spp. However, soil bacteria do not produce antibiotics at levels comparable to commercial applications – which can occasionally be in the scale of thousands of kilograms [13, 15, 16]. Also, the efficiency of superficial application of antibiotics in limiting the growth of plant bacterial pathogens, including some that are intracellular, is unclear. Recent studies have suggested injection of oxytetracycline produces better results [19, 29]. The spread of antibiotic resistance has been documented from agricultural use for antibiotics like tetracycline and streptomycin [30–32]. It has been noted that antibiotic resistance genes are naturally found in the environment [33, 34]. Therefore, application of consistent selection pressure by excessive and frequent use of antibiotics may enrich the population of naturally resistant organisms. However, at least in some instances under certain conditions, it was noted that streptomycin use did not alter the composition of soil microbial communities appreciably [35, 36].

Several reports on degradation kinetics and mechanisms of degradation of the antibiotics that are discussed here are available [21, 37–44]. It has been reported that the half-life of oxytetracycline at 25 °C is approximately 7 days, at 35 °C is 3 days and at 60 °C is 0.2 day, indicating a rapid temperature-dependent degradation of oxytetracycline, as the half-life at 4 °C is 120 days [37]. According to the same study, the half-life due to photolysis in the presence of sunlight is in the same order of magnitude. A similar investigation exists evaluating the photostability and temperature stability of streptomycin [44]. Briefly, the photodegradation of streptomycin is more modest than oxytetracycline by nearly 10-fold. The half-life of streptomycin was determined to be nearly 105, 42 and 30 days at 15 °C, 25 °C, and 40 °C respectively, implying a decreased rate of degradation when compared to oxytetracycline. A description of the possible degradation products of oxytetracycline and streptomycin are available [37, 44]. Our results showing a faster loss of efficacy for oxytetracycline than streptomycin upon sunlight exposure are therefore in agreement with the reported degradation kinetics of these antibiotics. To our knowledge, analysis such as the one we have conducted to monitor the biological efficacy of antibiotics subsequent to exposure to environmental elements are either lacking or not publicly available (as recognized by this article [14]). Our experimental conditions simulate the agricultural use of antibiotics and our results indicate that sunlight (heat and/or ultraviolet radiation) contributes to the degradation of oxytetracycline and streptomycin. Although our report is limited in scope, we believe it sheds light on the fate of antibiotics in the environment. Further studies to understand the effects of antibiotics are needed to inform the public and appropriate regulatory agencies [2–4].

## MATERIALS AND METHODS

### Strains used and general methods

The *B. subtilis* strain PY79 and the *E. coli* strain K-12 were incubated in 2 ml LB at 37 °C and grown until the culture OD_600_ reached 1.0 (exponential growth phase). A 100 μl aliquot of culture was then spread onto LB agar plates using sterile beads and set to dry completely prior to the placement of discs, see section below.

### Disc-diffusion assay

UV sterilized Whatman filter paper discs (7 mm) were impregnated with 5 μl of a freshly made stock antibiotic solution of either 40 mg/ml streptomycin sulfate (MilliporeSigma) in sterile distilled water or 10 mg/ml oxytetracycline hydrochloride (Alfa Aesar) in sterile distilled water to reach a concentration of 200 μg for streptomycin and 50 μg for oxytetracycline in each disc, and then set to dry completely. The concentrations selected were based on the concentration range recommended for agricultural use [45], and after empirically ensuring similar initial zones of inhibition for both antibiotics in the strains tested. To mimic the use of agricultural antibiotics, the discs were then placed outdoors (during spring months in Tampa, FL, USA where the average daytime temperature ranged from 27 to 32 °C) in direct sunlight for 7 or 14 consecutive 24-h periods (days) in parafilm-sealed sterile Petri dishes. Discs that were kept indoors in a dark cabinet at room temperature for 7 or 14 days, freshly prepared discs made the day of testing, and 5 μl of sterile water were used as controls. Discs were then transferred and pressed onto the pre-inoculated LB agar plates and incubated overnight at 37 °C. The zone of inhibition measurements were taken from the center of the disc to the edge of the zone of inhibition, minus disc radius (3.5 mm).

### Statistical analysis

GraphPad Prism Software (version 8.3.1) was used to analyze the data. All data represent biological triplicate data with technical replicates. Graphs show mean values and error bars represent standard deviation (SD).

## ACKNOWLEDGEMENTS

We thank our lab members for comments on the manuscript and assistance with data visualization. This work was funded by a start-up grant from USF (PE). A preprint of this manuscript is available on bioRxiv [46].

## AUTHOR CONTRIBUTIONS

The conception and design of the study (SK, PE), data acquisition (SK, AO), analysis and/or interpretation of the data (SK, AO, PE), and writing of the manuscript (SK, PE).

